# Growth Optimization Predicts Microbial Success in a Permafrost Thaw Experiment

**DOI:** 10.1101/2025.09.01.673550

**Authors:** JL Weissman, Joy M. O’Brien, Nathan D. Blais, Hannah Holland-Moritz, Katherine Shek, Robyn A. Barbato, Thomas A. Douglas, Jessica Gilman Ernakovich

## Abstract

Ongoing climate warming is thawing global permafrost, making vast pools of organic carbon available as microbial growth substrates. Uncertainty surrounding microbial successional dynamics limits our ability to parameterize the global-scale biogeochemical consequences of this thawing permafrost. We developed a genomic index of growth optimization to predict whether individual taxa increase or decrease in abundance during early thaw and validate our approach using incubation experiments from permafrost collected in Interior Alaska.

## Introduction

Permafrost soils, perennially-frozen soils that underly 15% of land area in the Northern Hemisphere [1], contain twice as much carbon as is currently in Earth’s atmosphere [2]. As ongoing and future climate warming thaws permafrost, these vast pools of organic carbon become available as growth substrates for microorganisms [3]. The resulting microbial feast will release carbon dioxide (CO_2_) and methane (CH_4_) into the atmosphere at rates on the same order of magnitude (between 55 – 232 Pg of CO_2_ equivalent from 2000-2099) as total US emissions [2]. Yet, how these microbial communities respond to thaw in the near- and medium-term, and, more specifically, what leads to an organism’s success during this process, are still largely open questions [4]. This inability to accurately parameterize permafrost microbial community dynamics represents a significant challenge for developing mechanistic models of the global-scale biogeochemical consequences of thawing permafrost [4].

Permafrost harbors taxonomically and metabolically diverse microbial communities, though many resident taxa are presumed to have low activity in intact permafrost [5]. Upon thaw, rapid changes in biophysical conditions lead to large shifts in microbial community taxonomic and functional composition [6, 7]. Predicting which microorganisms will proliferate immediately after thaw, especially during abrupt thaw such as in thermokarst formation [8], is a necessary first step in the process towards building microbially explicit process models to constrain the role of the permafrost-climate feedback in the Earth system. While the long-term successional dynamics of these communities will likely lead them to diverge considerably from their early-thaw composition following shifts in available carbon sources and redox states, priority effects in thawing permafrost systems mean that early dynamics can have persistent impacts on long-term community outcomes [9]. Key to predicting early-thaw microbial dynamics is the development of tools that can assess the competitive abilities of the post-thaw microbiome at the level of individual taxa.

Genomic traits associated with rapid growth can serve as useful indicators of organismal competitiveness during times of high resource abundance, such as those in recently thawed permafrost. Maximum growth rate, which can be estimated from genomes and metagenomes using signatures of translation optimization [10, 11], strongly predicts instantaneous growth dynamics across systems, making it a key parameter in population models. For example, experiments in marine systems have shown that genomically-derived estimates of microbial maximum growth rates are strongly correlated both with instantaneous growth rates inferred from incubation experiments [12] and instantaneous respiration rates resolved at the single-cell level from natural coastal communities [13]. In temperate soils, genomic estimates of maximum growth rate strongly correlated with bacterial growth and transcription after rewetting [14]. We hypothesize that early microbial dynamics in recently thawed permafrost will be determined largely by growth maxima, as bacteria take advantage of previously frozen organic carbon. Here, using controlled experiments, we show that genomic characteristics associated with translation optimization and high-resource “copiotrophic” lifestyles predict whether organisms will increase or decrease in relative abundance during early permafrost thaw.

## Results and Discussion

### Community-Wide Average Maximum Growth Rate Predictions Reveal a Shift in Growth Strategies During Thaw

We performed a 2 °C thaw experiment on permafrost sampled from three Alaskan field sites in a controlled laboratory environment (see Methods; [15]). We then sequenced metagenomes and applied the growth-rate prediction tool *gRodon* [11] to estimate the average community-wide maximum growth rate from metagenomes sampled pre- and post-thaw from our experiment. This tool leverages genomic signatures of translation optimization, primarily the codon usage bias of ribosomal proteins, to predict the maximum growth rate of an individual organism or community of organisms directly from a genome or metagenome sequence [11]. We hypothesized that the increase in bioavailable carbon post-thaw would favor a community comprised of fast-growing copiotrophs as compared to the pre-thaw microbial community in these experiments [3]. This expectation was borne out, where samples post-thaw nearly always had lower average predicted minimum doubling times (*i*.*e*., faster maximum growth rates) than their pre-thaw counterparts (Fig. 1a; two-tailed paired t-test, t=4.7, df=12, p=5.0e-4). These changes in community growth potential were often large, with a shift of 3.3 hours on average in the predicted average minimum doubling time across cores, with individual experiments showing shifts up to 8.1 hours in the predicted doubling times (Fig. 1b).

**Figure 1:**
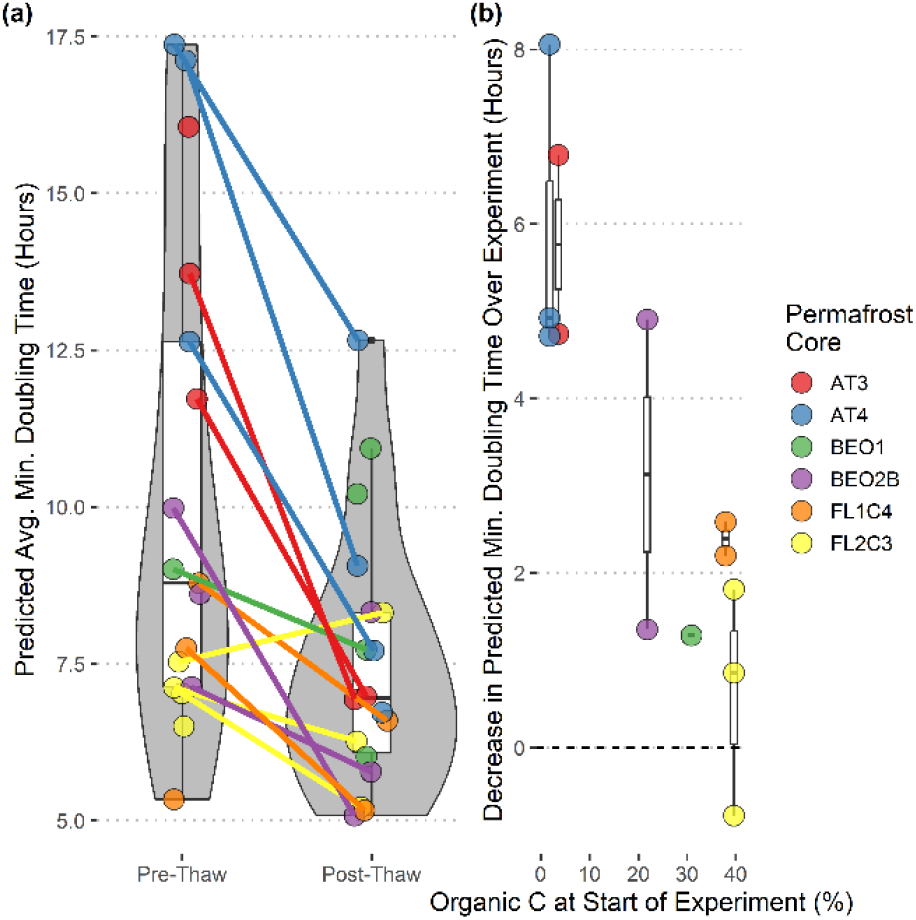
Metagenomic growth rate prediction shows clear shifts in community growth potential over the course of a controlled permafrost warming experiment. (a) The predicted average community-wide maximum growth rate (metagenome prediction mode in gRodon using all assembled contigs) shows a consistent shift in community-wide growth potential during thaw. Colored lines connect pre- and post-thaw samples for the same experimental replicates, for replicates where both samples had sufficient annotated ribosomal proteins for growth prediction. (b) The predicted change in community average maximum growth rate during the thaw experiment is negatively correlated with the percent organic soil carbon in each sample at the start of the incubation.

We hypothesized that cores with a higher percentage organic carbon by mass would see the growth potential of their microbial communities shift more dramatically during the thaw process. Surprisingly, we found the opposite result, that low-carbon samples saw the greatest increase in community maximum growth rates (Fig 1b; Pearson’s product-moment correlation, rho=0.84, t=5.2, df=11, p=3.1e-4), leading to an overall lower variability in predicted community growth rates among post-thaw samples than pre-thaw samples. Though, as expected, high-carbon samples had faster community maximum growth rates than the low-carbon samples both pre- and post-thaw (S1 Fig.).

### Metagenome-Assembled Genomes Cluster Along an Axis of Copiotrophy that Predicts Changes in Relative Abundance During Thaw

We reconstructed 369 genomic bins from pre- and post-thaw metagenomes. Of these, 102 had sufficiently low contamination (<5%) and a high enough number of annotated ribosomal proteins (≥10) to perform maximum growth rate prediction (Fig S2; [11]). Bins reconstructed from post-thaw contigs had generally faster inferred growth rates than those from pre-thaw contigs (Fig S2; two-sided Welch’s t-test, t=5.3, df=87, p=9.0e-7) This bin-level pattern reflects community-wide patterns (Fig 1a) though these bins only accounted for a small fraction of the overall community (7% of total relative abundance on average). Overall, bins represented a wide range of predicted growth strategies, with predicted minimum doubling times at a reference temperature of 2 °C ranging from 3.1 to 41.1 hours (Fig 1d).

Recent work in marine systems developed an “index of copiotrophy” combining key traits related to growth strategy (codon usage bias of the ribosomal proteins, genome size, and the number of carbohydrate active enzymes) to quantify the growth strategies of microbes, with distinct biogeographic patterns linked to carbon supply and lability separating along this single axis of variation [16]. Codon usage bias measures translation optimization and the main input into our genomic growth predictions. Both genome size and carbohydrate-active enzymes are indicators of adaptation for a metabolically flexible lifestyle associated with high-resource environments. Here, we took the first component of a principal component analysis of these three traits across our 102 bins as our index of copiotrophy in these permafrost samples. Along this axis we defined two distinct clusters of bins, which we will refer to as the “oligotrophic” and “copiotrophic” clusters, as well as a region of hard-to-define bins between these clusters (where classification uncertainty ≥10%), using a Gaussian-mixture model (Fig 2a-b). While definitions of copiotrophy and oligotrophy vary in the literature, here we use our previously proposed evolutionary definition in which fast-growing “copiotrophs” are genomically optimized for high-resource environments and slow-growing “oligotrophs” are optimized for low-resource environments [11, 17, 18]. Importantly, the definition we use here is not based on the lability of preferred carbon sources, consistent with the definition used by marine scientists (where prototypical oligotrophs specialize on highly-labile carbon sources at low concentration [16]), but contrasting with common definitions in soil microbiology. Our clusters showed phylogenetic affiliation, with most copiotrophic bins appearing in the Proteobacteria and Actinobacteria (though these phyla had a wide range of predicted growth strategies), while other phyla such as the Acidobacteriota and Dormibacterota were primarily oligotrophic (Fig 2c). These differences broadly align with the expected roles of these groups in soil ecosystems [19, 20], with the latter groups being particularly challenging to cultivate in the laboratory.

**Figure 2:**
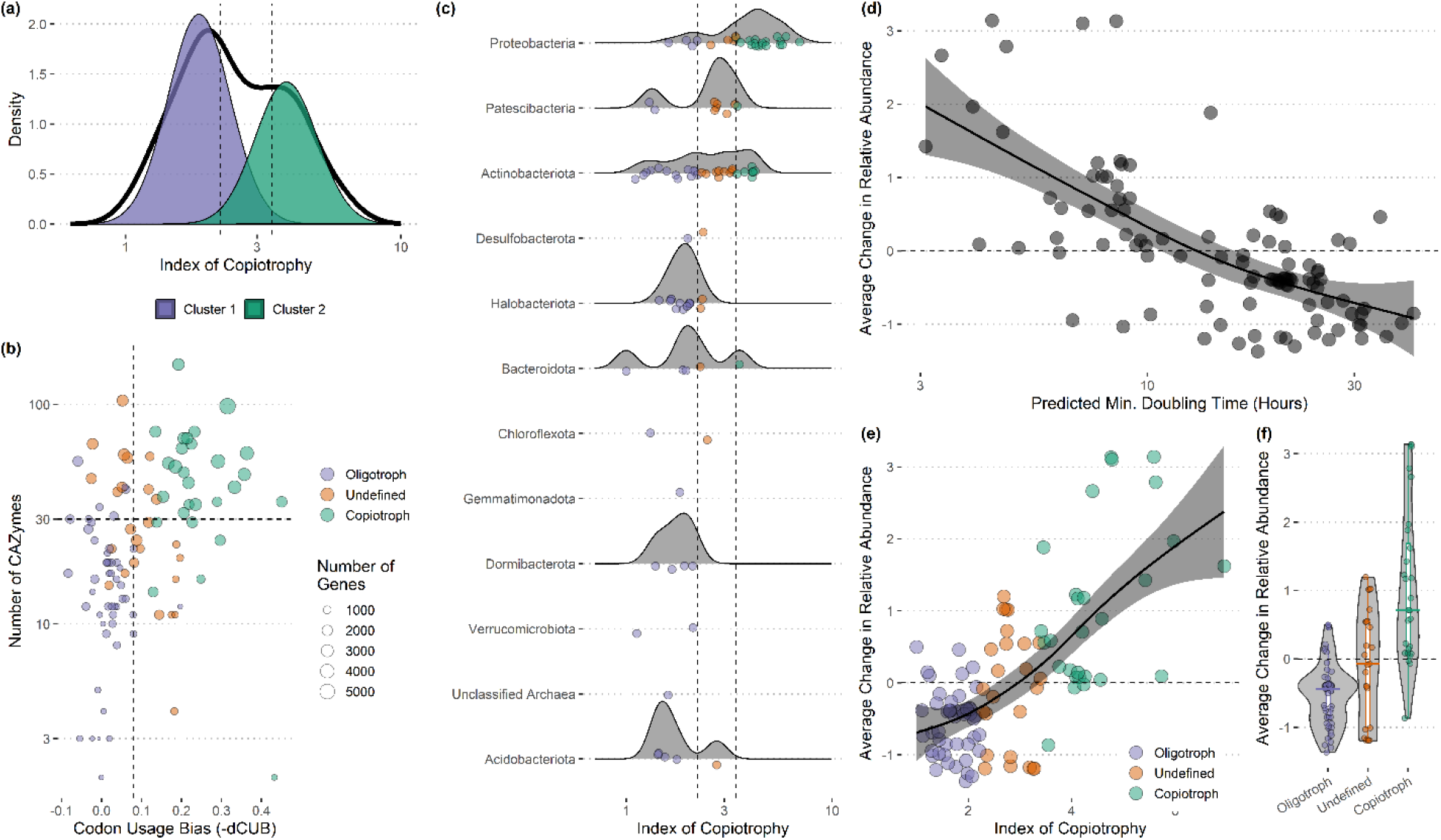
An index of copiotrophy predicts organismal (bin-level) success during a permafrost thaw experiment. (a) Bins cluster into two groups of high and low copiotrophy index. Vertical dashed lines surround region where classification uncertainty increases above 10% (bins in this region classified as “undefined”). (b) The three genomic variables in the copiotrophy index are highly correlated across bins (codon usage bias, number of CAZymes, and total number of genes for each bin). Vertical dashed line at cutoff generally used to separate copiotrophs and oligotrophs on the basis of codon usage bias (-dCUB=0.08). Horizontal dashed line at 30 denotes approximate divide between copiotrophs and oligotrophs as defined by the index of copiotrophy. (c) Phyla vary widely in their index of copiotrophy. Points show individual bins. Vertical lines as in (a). (d) The change in transformed relative abundance (difference in post- and pre-thaw transformed relative abundance of each bin) of a bin decreases with increasing genomically-predicted minimum doubling time. (e-f) The change in transformed relative abundance of a bin increases with increasing index of copiotrophy, with most copiotrophic bins increasing in abundance pre-to post thaw and most oligotrophic bins decreasing in abundance.

We hypothesized that genomic indicators of a copiotrophic lifestyle, such as a fast predicted maximum growth rate and a high index of copiotrophy, should be positively associated with an organism’s fitness during permafrost thaw, since during the initial stages of thaw previously-unavailable sources of organic carbon flood the system to create a resource-rich environment [3]. As our proxy for fitness, we took the change in relative abundance across pre- and post-thaw sample pairs for each bin. Indeed, these genomic characteristics appeared to predict success, with predicted minimum doubling time being negatively correlated with the average change in relative abundance of a bin across pre/post-thaw sample pairs (Pearson’s product-moment correlation, rho=-0.62, t=-8.0, df=100, p=2.7e-12; Fig 2d) and the index of copiotrophy being positively associated with a bin’s average change in relative abundance (Pearson’s product-moment correlation, rho=0.68, t=9.2, df=100, p=6.5e-15; Fig 2e-f). Strikingly, 25 out of 28 copiotrophic bins showed increases in relative abundance whereas 43 out of 49 oligotrophic bins showed decreases in relative abundance, with undefined bins showing a nearly even split (13 out of 25).

In conclusion, our index of copiotrophy appears to be a useful metric for predicting whether a population will increase or decrease in relative abundance during the initial stages of community assembly during permafrost thaw (Fig 2e-f). Simple functional metrics like our index that can be derived directly from genomic characteristics further the development of metagenomics into a predictive, rather than a simply descriptive, tool for understanding the trajectories of microbiomes in a changing climate.

## Methods

### Sample Collection and Incubations

Detailed sample collection and incubation protocols are described in [15]. Briefly, controlled thaw experiments were performed on permafrost cores taken from three permafrost sites in interior and northern Alaska. These include a site above the Cold Regions Research and Engineering (CRREL) Permafrost Tunnel (AT), the CRREL Farmers Loop Experimental Station (FL), and the Barrow Experimental Observatory (BEO). All three sites comprise carbon- and ice-rich permafrost that is often susceptible to thermokarst formation upon thaw (see [15] for further site and sampling details). Each site had two cores, from which we subsampled four replicates. Frozen soil cores were gradually thawed at 2 °C for 96 days. Total carbon from dry soil was measured via loss on ignition at a combustion temperature of 950C (Costech ESC 4010, Valencia, CA USA), instrument standards (acetanilide) were run every 12 samples, and R^2^ was >0.9998 across the whole range.

### DNA Extraction and Sequencing

DNA was extracted from 0.25 g of soil using the Qiagen PowerSoil Pro kit (Qiagen, Hilden Germany), following manufacturer’s protocol. Samples were prepped with Illumina’s TruSeq DNA Library Prep Kit v2. They were quantified with a qubit Qubit HS Assay, for Invitrogen’s Qubit Fluorometer (Catalog number: Q32851 or Q32854) and purified with Blue Pippin Prep DNA Gel Cassettes (1.5%; catalog number: BDF3010). Samples were run on a 2X151 P1 Nextseq run at the Next Generation Sequencing Core Facility at Argonne National Laboratory (Lemont, IL, USA).

### Sequence Analysis

Adapters and low-quality reads were trimmed using fastp v0.23.4 with default settings [21]. Reads from each sample were assembled using the SPAdes v4.0.0 genome assembler with option “--meta” (metaSPAdes; [22]). Coverage of each contig across all samples was calculated using fairy v0.5.7 [23]. Metagenomic bins were then inferred from bins for each sample, using coverages across all samples, with MetaBAT2 v2.17 with a minimum contig length set to 1.5kb [24]. Bin quality was assessed using CheckM2 v1.0.1 [25].

Bins and metagenomic contigs were annotated with prokka v1.14.6 [26] and dbcan v 4.1.4 [27]. We predicted the maximum growth rate of each bin using gRodon v 2.4.0, as well as for each mixed-species metagenome, using metagenome mode version 2 [11, 28]. Using gRodon’s metagenome mode, we predict an average maximum growth rate across an entire community by considering all genes called from all assembled contigs (not only binned contigs) and weighting the average by the coverage of each gene. This captures an average that includes organisms that cannot be binned [28]. For gRodon estimates where there were less than 10 predicted ribosomal proteins in the source genome used for prediction, the resulting estimates were discarded based on established cutoffs for this tool. Taxonomy was assigned to each bin using gtdbtk v2.1.1 [29]

To construct our index of copiotrophy across our bins we first took the first principal component of a PCA with of three variables (1) dCUB, (2) the total number of genes in a bin, and (3) the proportion of genes annotated as CAZymes. We then transformed these values to range from 1 to infinity by subtracting off the minimum value (which was negative) and adding one. This procedure is consistent with previous construction of the same metric for the analysis of ocean transects [30]. Clustering along this axis of variation was performed using a Gaussian-mixture model using the R package mclust v6.1.1 with default settings [31].

The relative abundance of each bin across metagenomic samples was assessed using coverM v0.7.0 [], and relative abundances were then transformed using the modified centered log-ratio transform (mclr) from the SPRING R package v1.0.4 [32]. This transformation effectively mitigates the impact of the fact that population data comes in the form of relative abundances on downstream statistics while also not requiring the addition of pseudocounts for zero abundance data cells [32].

## Data and Code Availability

Sequenced metagenomes are available under NCBI BioProject PRJNA1313352. Metagenomic bins and associated metadata are available under Zenodo DOI 10.5281/zenodo.17025131. Associated code is available at https://github.com/jlw-ecoevo/permafrost_warming.

---

**Supplemental Figure S1:**
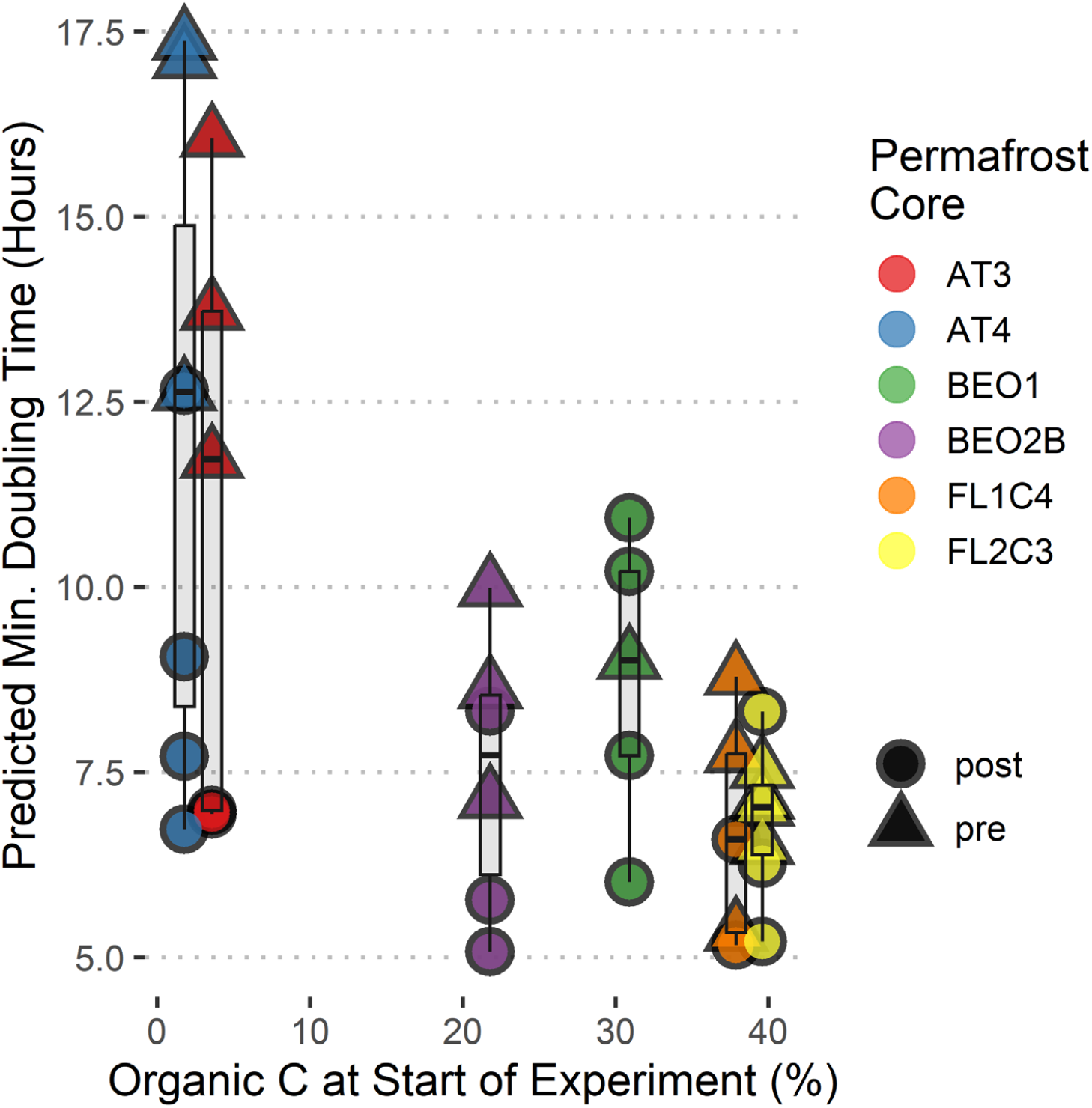
Predicted average community-wide minimum doubling times and organic carbon for each sample.

**Supplemental Figure S2:**
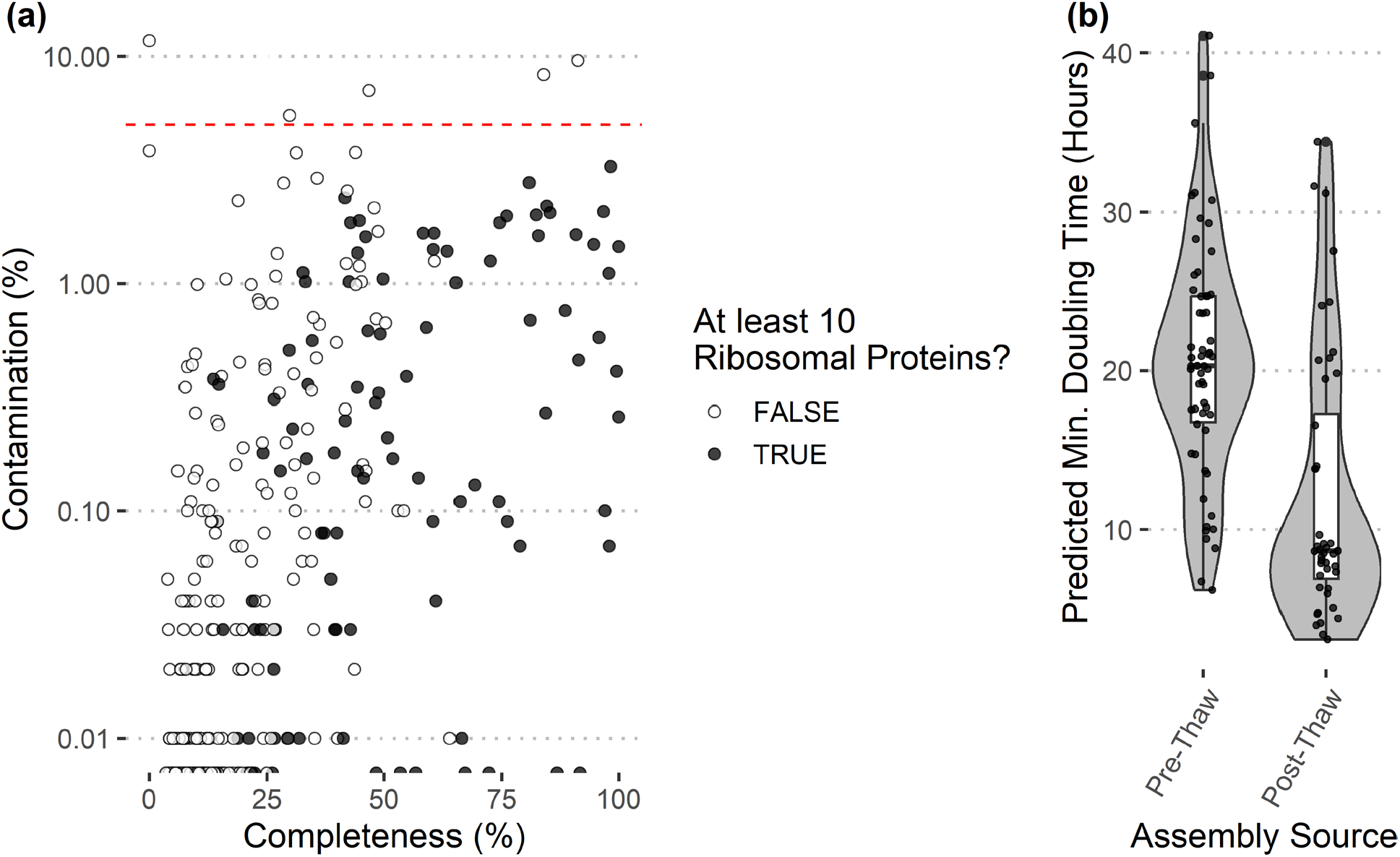
(a) Quality and contamination summary of bins. Bins used for growth rate prediction had at least 10 annotated ribosomal proteins and less than 5% contamination. (b) Maximum growth rate prediction from reconstructed metagenomic bins (genome prediction mode in gRodon) reveals that bins binned from pre-thaw contigs had longer minimum doubling times than those binned from post-thaw contigs.

## Notes

https://github.com/jlw-ecoevo/permafrost_warming

https://zenodo.org/records/17025132?token=eyJhbGciOiJIUzUxMiJ9.eyJpZCI6IjllNzUxMmE4LWYwNDEtNGJkNS1iMGVlLTA0ZDBiNDJhOTY3MSIsImRhdGEiOnt9LCJyYW5kb20iOiIzNTFmNzg2MTljNzJjYzJkMzgzZTU0MzRiMzcwNjJjMiJ9.trszfviP8LaNyJ5c8NDuPnanXF_ifLazTxzd1266AZaOymbL8KkblM5hSqGcKFFa0mZUDreHlzasdUwglZUH6g

https://www.ncbi.nlm.nih.gov/bioproject/?term=PRJNA1313352

### Competing Interest Statement

The authors have declared no competing interest.

### Summary of Updates

Added ORCID for an author and modified funding acknowledgement

### Funding

- National Science Foundation: #2144961
- UNH Collaborative Research Excellence Initiative
- NH NASA Space Grant Consortium
- DoD SMART Scholarship
- Applied Research Funding for Extreme Cold Weather through the ASA (ALT) and AFC

